# Refinement of the Sugar Puckering Torsion Potential in the AMBER DNA Force Field

**DOI:** 10.1101/2024.08.21.609021

**Authors:** Marie Zgarbová, Jiří Šponer, Petr Jurečka

## Abstract

The transition from B-DNA to A-DNA occurs in many protein-DNA interactions or in DNA/RNA hybrid duplexes, and thus plays a role in many important biomolecular processes that convey the biological function of DNA. However, the stability of A-DNA is severely underestimated in current AMBER force fields such as OL15, OL21 or bsc1, potentially leading to unstable or deformed protein-DNA complexes. In this study, we refine the deoxyribose dihedral potential to increase the stability of the north (N) puckering present in A-DNA. The new parameters, termed OL24, model A/B equilibrium in B-DNA duplexes in water in good agreement with NMR experiment. They also improve the description of DNA/RNA hybrids and the transition of the DNA duplex to the A-form in concentrated ethanol solutions. These refinements significantly improve the modeling of protein-DNA complexes, increasing their structural stability and A-form population, while maintaining accurate representation of canonical B-DNA duplexes. Overall, the new parameters should allow more reliable modeling of the thermodynamic equilibrium between A- and B-DNA forms and the interactions of DNA with proteins.

## Introduction

### Importance of A/B equilibrium

The equilibrium between the canonical B-DNA form and the less stable A-DNA is crucial for many processes. One of the best understood roles is in the interactions of DNA with proteins. DNA often undergoes a partial (local) transition to A-DNA, which is important for instance for transcription factors that specifically bind to A-DNA prone sequences, such as the TATA-box.^1^ Many polymerases also rely on locally inducing the A-form in DNA strand^2, 3^ as do other proteins.^4-6^ Beyond protein-DNA interactions, the A/B equilibrium is important in DNA/RNA hybridization, thus affecting processes like RNA interference, telomere maintenance (telomerase RNA component/DNA hybrid) or CRISPR gene.^7, 8^ Accurate modeling of these molecules requires a reliable description of the A/B DNA conformational properties.

### The A and B Forms of DNA

The A-form of DNA differs from the B-form in several geometrical parameters, including the inclination of base pairs to the helical axis, x-displacement (which describes the “hole” in the axial view of A-DNA), slide, roll, and groove widths. A- and B-DNA forms are depicted in Figure 1, along with inclination and x-displacement and their distribution in X-ray structures (data taken from non-complexed A- and B-DNA from Ref. ^9^). In our previous work, we chose x-displacement and inclination (Figure 1) as suitable parameters for monitoring the B/A transition in MD simulations.^10^ However, when the A-DNA form is stable enough, the RMSD value relative to the A-form also serves as a good indicator.

**Figure 1.**
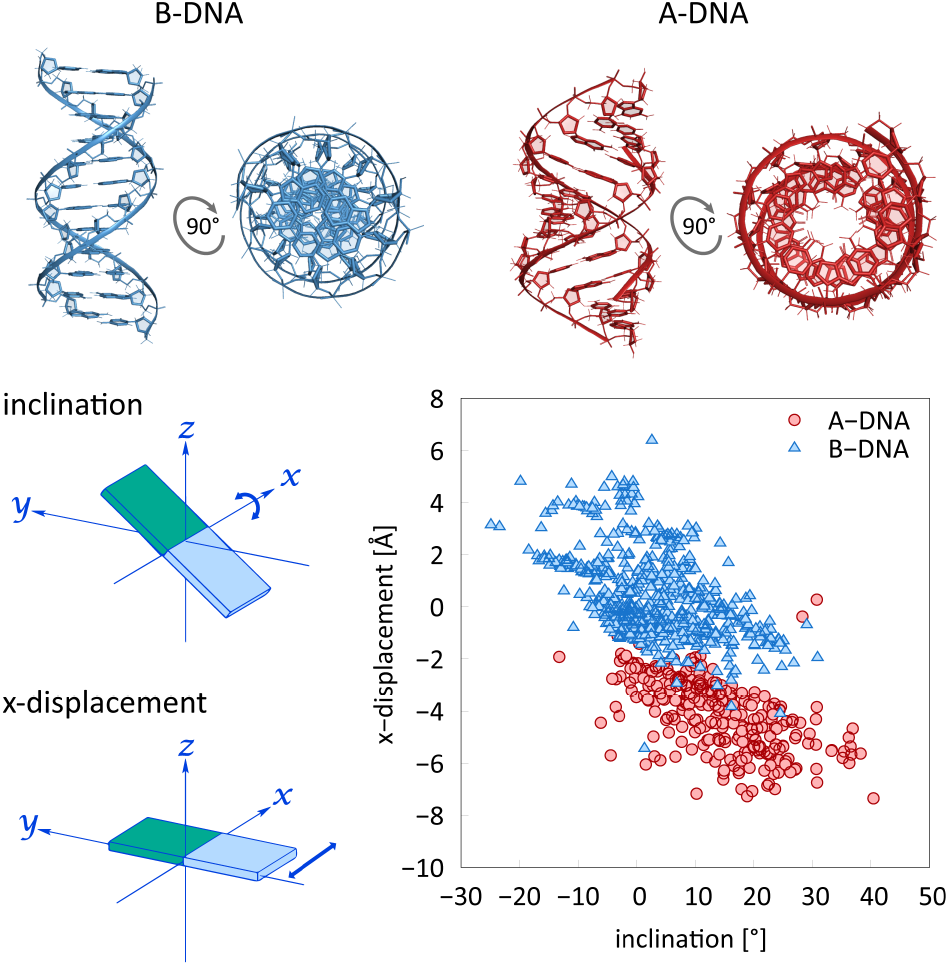
B-DNA and A-DNA. Inclination and x-displacement for B- and A-DNA forms, measured in X-ray data taken from non-complexed DNA double strands.^9^

The backbone conformations of the A- and B-forms also differ: A-DNA features a north (N) sugar puckering, while B-DNA has a south (S) puckering. Additionally, the glycosidic torsion angle χ is in the *anti* region (∼200°) for A-DNA and in the high-*anti* region (∼ 250°) for B-DNA. For definition of backbone dihedrals see Figure 2A. It is important to note that the B to A transition is not necessarily a two-state process from a point of view of a single nucleotide. Intermediate forms can exist where A-like puckering is combined with B-like χ and B-like puckering with A-like χ. These intermediate forms can be involved in protein-DNA interactions.^11^ In the following text, we will focus on the fraction of N deoxyribose puckering as the primary indicator of the A-form character. For the distribution of the pseudorotation angle (P) in non-complexed B-DNA and A-DNA X-ray structures^9^ see Figure 2B). Typically, the χ = anti accompanies the N puckering; when this is not the case, it will be explicitly noted.

**Figure 2.**
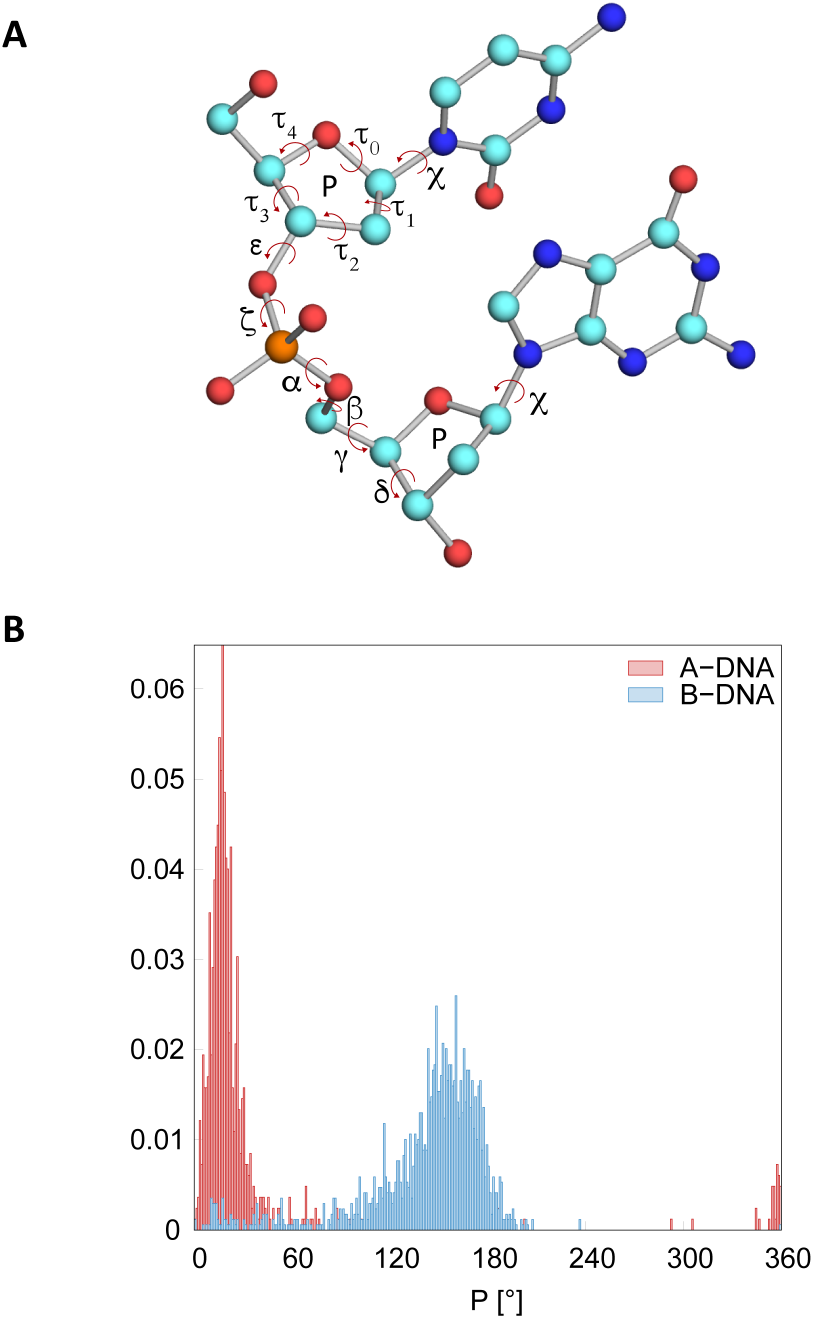
(A) Backbone dihedral angles and pseudorotation angle P. (B) Distribution of P in A-DNA and B-DNA molecules.

### Force Field Modeling of A-DNA

The modeling of the A/B equilibrium in AMBER force fields (*ff*) has been a long standing problem, even with recent refinements such as OL15,^12^ OL21,^13^ bsc0^14^ or bsc1^15^. All these *ff*s tend to be strongly biased towards the B-DNA form. This leads, e.g., to instability of the A-DNA form in the concentrated (85%) ethanol solutions, where A-DNA is experimentally observed.^16^ In molecular dynamics (MD) simulations, very fast spontaneous transition from A-DNA to B-DNA has been observed in concentrated ethanol solution for bsc0 and similarly for bsc1 and OL15,^10^ and OL21 as reported herein. Inconsistency with experimental data has also been observed in simulations of the Dickerson-Drew dodecamer (DDD) in aqueous solution, which should populate approximately 12-15% of the N sugar puckering according to NMR experiment.^17, 18^ However, bsc0, bsc1 and OL15 show only marginal sampling of N puckering.^10^ This underestimation of N puckering is also evident in other DNA duplexes and free nucleosides in aqueous solution.^10^ Recent studies have also highlighted a systematic underestimation of the A-form in MD simulations of protein-DNA complexes^11^ and a very low content of N puckering in DNA/RNA hybrids.^19^ In contrast, the AMOEBA^20, 21^ *ff* shows a significantly higher population of N puckering in DNA/RNA hybrids and CHARMM36^22^ shows perhaps too high a population^19^ (see also DRUDE-2017^23, 24^ results for DNA duplexes^25^). These observations indicate an unsatisfactory description of the A/B equilibrium in current AMBER *ff*s.

The instability of the A-DNA form can have significant consequences in MD simulations. For instance, in protein-DNA complexes, the DNA often adopts the A-form conformation. However, in simulations using the mentioned AMBER *ff*s, the A-form can rapidly disappear, leading to deviations from the initial X-ray structure over time, as evidenced for example by the increased root mean square deviation (RMSD).^11^ The large DNA geometrical changes inevitably obscure the interpretation of the simulations. Furthermore, even when the geometry of DNA remains unchanged on shorter time scales, inaccurate modeling of the A/B equilibrium can still pose a problem. The transition from the A-DNA backbone state to the B state is accompanied by a free energy change of approximately 1 kcal/mol per nucleotide.^16^ Inaccurate modeling of this transition in simulations can obviously bias the stability of the protein-DNA complexes, leading to incorrect predictions of association Gibbs energies. In some cases, protein-DNA complexes have been observed to dissociate in simulations over longer time scales.^26^ These issues raise concerns about the reliability of modeling protein-DNA interactions using current AMBER *ff*s.

The transition to the A-form is also important for DNA/RNA hybridization because the DNA strand in the hybrids exhibits a higher content of A-like sugar puckering.^27^ Again, even if the geometry of the hybrid geometry is modeled reasonably well, the free energy of hybridization is likely to be underestimated if the transition towards the A-form in DNA is not correctly modeled.

### OL24 Force Field Refinement

We have developed a modification of the OL21 AMBER *ff*, named OL24, which increases stability of the A-DNA form. Key modifications include the δ sugar phosphate backbone torsion and the τ1 deoxyribose ring torsion parameters. Additionally, we have adopted the glycosidic torsion potential from the OL3 RNA *ff*,^28^ replacing the glycosidic torsion potential used in OL15 and OL21 DNA *ff*s, which was specifically derived for DNA.^29^ Thus, both RNA and DNA OL *ff*s now share the same glycosidic torsion.

The δ and τ1 modifications are based on fitting quantum mechanical (QM) data using our methodology, which accounts for conformation-dependent solvation effects while avoiding double counting of solvation energy.^30^ The OL24 parameterization retains the very good performance of OL15 and OL21 *ff*s^31, 32^ for modeling the B-DNA double helices. Importantly, it provides a better representation of A-type nucleotide populations, more consistent with NMR experiments for DNA duplexes in aqueous solution. OL24 models spontaneous transition to A-DNA in concentrated ethanol solutions, and significantly improves the description of protein-DNA complexes and DNA/RNA hybrids. In the following, the OL24 simulations are primarily compared with the previous OL21 Olomouc *ff* variant. We do not report the results of the older OL15 and bsc1 *ff* variants, as they are quite similar to OL21 for B-DNA and A-DNA duplexes and have been detailed in our previous works.^10, 11, 13^

## Methods

### Parameter development

The parameters for the deoxyribose pucker were derived by fitting reference QM dihedrals using a previously published methodology, in which the molecule is optimized at both QM and MM levels, with solvation effects included in both QM and MM calculations.^30^ For the QM calculations, the COSMO solvation model^33^ was used (PBE/QZVPP^34, 35^ level) and the PB model^36, 37^ was employed for the MM part (water, ε_r_ = 78). QM calculations begun with a constrained PBE/QZVPP optimization, followed by single-point energy calculations at a higher level of theory. The CCSD(T) complete basis set estimate was obtained by extrapolating MP2 energy using aug-cc-pVTZ and aug-cc-pVQZ basis sets and separate extrapolation of the HF and correlation energy,^38, 39^ and then adding a correction for higher order correlation effects calculated at the CCSD(T)/aug-cc-pVDZ level.^40^ The solvation energy calculated at the PBE/QZVPP level was added to the single point-energy. The model compound used was the abasic deoxyribonucleoside, 1,2-dideoxy-D-ribofuranose. In the pseudorotation angle scan, the dihedral angles τ1 and δ were constrained to values corresponding to given P values (0-360° at 10° intervals). The structure was optimized using TurboMole 7.3 software.^41^

### MD simulations

The MD simulations started from X-ray structures with PDB IDs 1BNA^42^ (DDD), 3VJV^43^ and 3PVI^44^ (protein-DNA complexes), NMR structures 1DRR and 1RRD^27^ or from models built by NAB module of Amber^45, 46^ (C_4_G_4_,^47^ G_4_C_4_^48^ and d(CATTTGCATC) by Weisz et al.^49^). An overview of the simulations is given in Table 1.

**Table 1.** Overview of MD simulations. DDD-r stands for DDD duplex with restrained terminal base pairs, *ff*s for hybrids are in the order of DNA/RNA.

| Molecule | Sequence | *ff* | Time [μs] | # replicas |
| --- | --- | --- | --- | --- |
| B-DNA | DDD | OL24 | 5 | 1 |
| B-DNA | DDD-r | OL24 | 5 | 1 |
| B-DNA | C<sub>4</sub>G<sub>4</sub> | OL21 | 2 | 1 |
| B-DNA | C<sub>4</sub>G<sub>4</sub> | OL24 | 2 | 1 |
| B-DNA | G<sub>4</sub>C<sub>4</sub> | OL21 | 2 | 1 |
| B-DNA | G<sub>4</sub>C<sub>4</sub> | OL24 | 2 | 1 |
| B-DNA | Weisz | OL21 | 2 | 1 |
| B-DNA | Weisz | OL24 | 2 | 1 |
| DNA/RNA hybrid | 1DRR | OL21/OL3 | 2 | 1 |
| DNA/RNA hybrid | 1DRR | OL24/OL3 | 2 | 1 |
| DNA/RNA hybrid | 1RRD | OL21/OL3 | 2 | 1 |
| DNA/RNA hybrid | 1RRD | OL24/OL3 | 2 | 1 |
| protein/DNA | 2VJV | ff14SB/OL21 | 1 | 3 |
| protein/DNA | 2VJV | ff14SB/OL24 | 1 | 3 |
| protein/DNA | 3PVI | ff14SB/OL21 | 1 | 3 |
| protein/DNA | 3PVI | ff14SB/OL24 | 1 | 3 |
| B-DNA/EtOH | DDD | OL21 | 1 | 3 |
| B-DNA/EtOH | DDD | OL24 | 1 | 3 |
| Nucleoside | DCN | OL21 | 5 | 1 |
| Nucleoside | DCN | OL24 | 5 | 1 |
| Nucleoside | DTN | OL21 | 5 | 1 |
| Nucleoside | DTN | OL24 | 5 | 1 |
| Nucleoside | DAN | OL21 | 5 | 1 |
| Nucleoside | DAN | OL24 | 5 | 1 |
| Nucleoside | DGN | OL21 | 5 | 1 |
| Nucleoside | DGN | OL24 | 5 | 1 |

The DNA, RNA and protein *ff*s used are detailed in Table 2. For DNA duplexes, hybrids, and protein-DNA complexes in aqueous solution, the starting structure was first neutralized with K^+^ ions and, the ion concentration was then adjusted to 0.15 M using KCl with parameters of Joung and Cheatham.^50, 51^ For simulations in 85% ethanol solution, the DNA was only neutralized with Na+ cations.^50, 51^ No ions were used in nucleoside simulations. SPC/E water model^52^ was used in all simulations except for water/ethanol mixtures, where TIP3P^53^ was used for compatibility with the OPLS ethanol parameters.^54^ A rectangular box with a 12 Å buffer was used for C_4_G_4_ and G_4_C_4_ simulations, a rectangular box of 60 × 85 × 85 Å for DDD simulations in ethanol/water and an octahedral box with a 12 Å buffer for the remaining simulations.

**Table 2.**
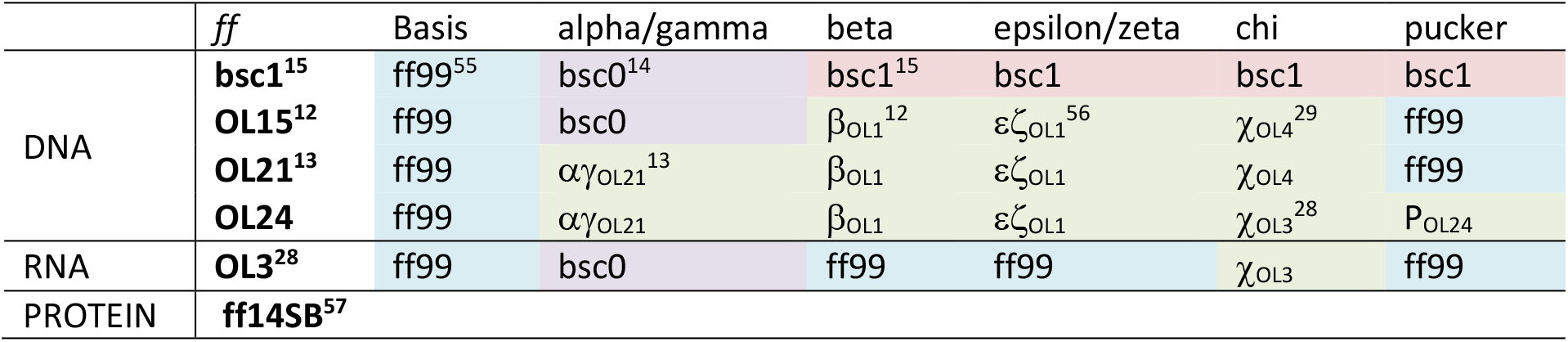
The DNA, RNA and protein *ff*s used. All dihedral angle modifications are based on ff99 *ff*.

|  | *ff* | Basis | alpha/gamma | beta | epsilon/zeta | chi | pucker |
| --- | --- | --- | --- | --- | --- | --- | --- |
| DNA | **bsc1**<sup>15</sup> | ff99<sup>55</sup> | bsc0<sup>14</sup> | bsc1<sup>15</sup> | bsc1 | bsc1 | bsc1 |
| | **OL15**<sup>12</sup> | ff99 | bsc0 | $\beta_{OL1}$<sup>12</sup> | $\epsilon\zeta_{OL1}$<sup>56</sup> | $\chi_{OL4}$<sup>29</sup> | ff99 |
| | **OL21**<sup>13</sup> | ff99 | $\alpha\gamma_{OL21}$<sup>13</sup> | $\beta_{OL1}$ | $\epsilon\zeta_{OL1}$ | $\chi_{OL4}$ | ff99 |
| | **OL24** | ff99 | $\alpha\gamma_{OL21}$ | $\beta_{OL1}$ | $\epsilon\zeta_{OL1}$ | $\chi_{OL3}$<sup>28</sup> | P<sub>OL24</sub> |
| RNA | **OL3**<sup>28</sup> | ff99 | bsc0 | ff99 | ff99 | $\chi_{OL3}$ | ff99 |
| PROTEIN | **ff14SB**<sup>57</sup> |  |  |  |  |  |  |

Initial relaxation was performed using a multi-step protocol described earlier.^58^ For DDD simulations in ethanol/water solution, the relaxation step in which the DNA was restrained and the solution with ions was allowed to move was extended from 10 ps to 20 ns.

MD simulations were carried out in the PMEMD CUDA code of the AMBER 20 software suit^45, 59^ under NpT conditions (1 bar, 298 K) with Langevin thermostat (gamma_ln = 5), Monte Carlo barostat (taup = 2), hydrogen mass repartitioning^60, 61^ and 4 fs time step, 10 Å direct space cutoff and SHAKE constraints on bonds involving hydrogen atoms with a default tolerance (0.00001 Å). The nonbonded pair list was updated every 25 steps. Coordinates of nucleic acids, proteins and ions were stored every 10 ps. For DDD simulations, additional runs with flat well restrains on the electronegative atoms of terminal base pair hydrogen bonds (flat between 2.5 and 3.2 Å, parabolic restraints with k = 20 kcal/(mol.Å^2^) for shorter and 30 kcal/(mol.Å^2^) for longer distances, respectively) were performed to allow a more accurate comparison of B-DNA helical parameters, minimizing the effect of fraying (denoted as “DDD-r” in Table 1).^58^

Free nucleosides were simulated with the χ angle restrained to the *anti* region using a flat well potential with a quadratic penalty of 10 kcal/rad^2^ applied below 160° and above 310° to prevent bias from intramolecular hydrogen bond, as described in Ref.^10^. The potential of mean force (PMF) was calculated from the probability distribution of the pseudorotation angle P in 3° bins using the formula PMF = −RT ln(nbin/nmax), where nbin is the count in each bin and nmax is the count of the most populated bin. The 95% confidence intervals for the counts in each bin (nbin) were calculated as ±1.96*SEM (SEM is standard error of the mean).

The analysis of DNA structural parameters was performed using the cpptraj module and the nastruct tool of the AMBER software package. Groove widths were calculated according to the method described by El Hassan and Calladine.^62^ For determining the north (N) sugar pucker populations, the pucker was classified as N if it fell within the interval −90° < P < 90°, and as south (S) otherwise.

Data presented in Figures 1 and 2 are calculated from a set of non-complexed DNAs^9^ and analyzed by X3DNA,^63^ excluding outliers (3σ). Illustrations of helical parameters are adapted from Ref.^64^

## Results and Discussion

### Parameter development

In OL24, potentials of the glycosidic angle χ and two dihedral angles in the deoxyribose ring were modified compared to OL15 and OL21. The χ_OL4_ parameterization, used in the previous OL15 and OL21 *ff* versions and originally developed for DNA,^29^ has been replaced by the χ_OL3_ parameterization from the RNA *ff*.^28^ This change was made because, in a previous study, χ_OL3_ was found to better stabilize the P/χ = A/A and B/A conformations in protein-DNA complexes.^11^ We show here that the χ_OL3_ parameterization does not adversely affect the simulations of the DNA molecules studied.

The two deoxyribose dihedral angles were initially refined based on high-level QM calculations. However, subsequent simulations showed that the fit to QM data still underestimated the stability of the N sugar pucker. Thus, the N minimum around P ∼ 18° was further stabilized to bring the %N into closer agreement with NMR experiments for DDD.^17, 18^ The PMF for the resulting OL24 parameters is shown in Figure 3.

**Figure 3.**
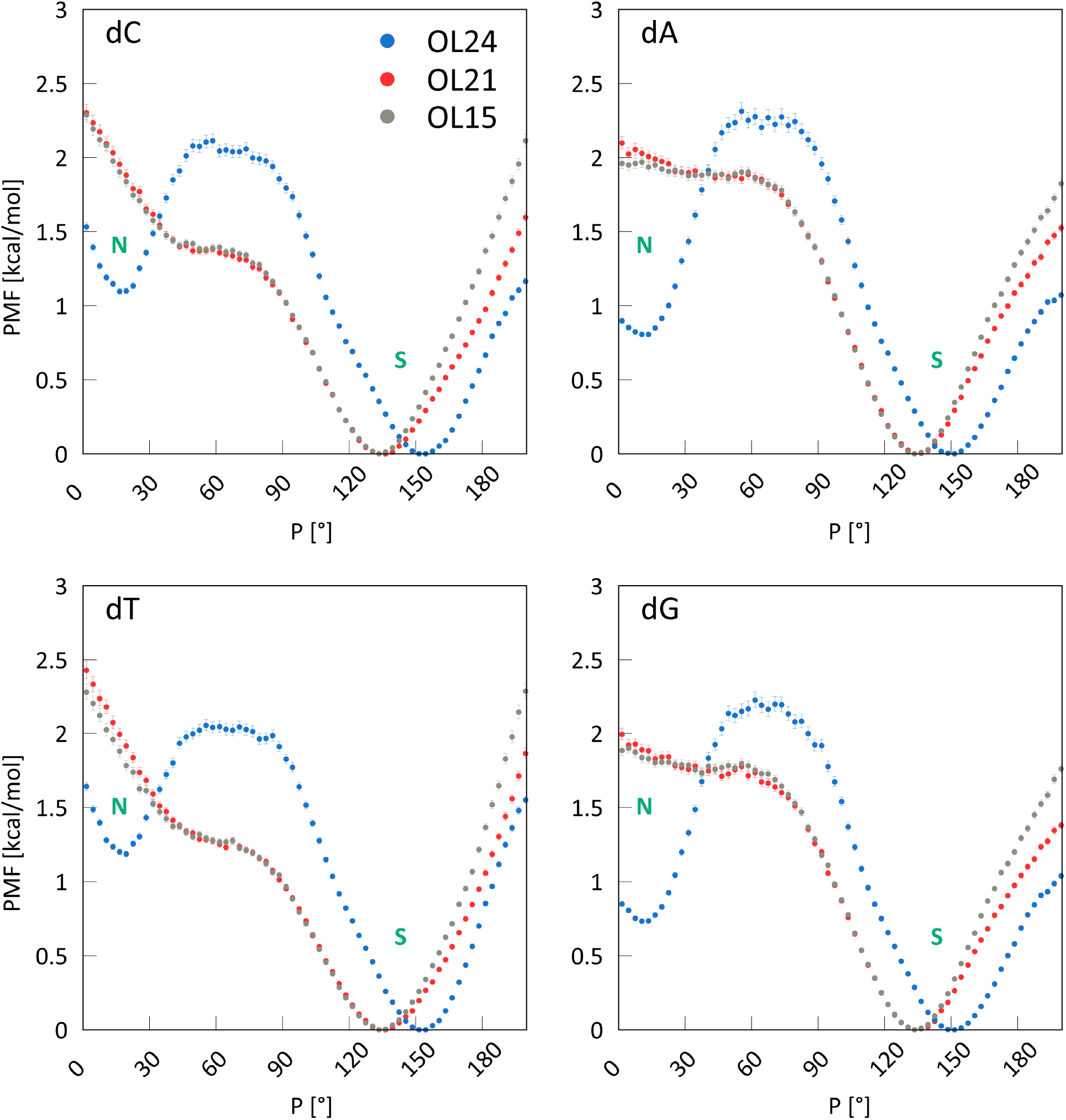
Potentials of mean force for DNA nucleosides calculated using OL15, OL21 and OL24 *ff*s in SPC/E water. The OL24 *ff* stabilizes the N region for all nucleosides.

The PMFs shown in Figure 3 for the OL15 and OL21 *ff*s are nearly identical, as they use the same χ and sugar pucker parameterization. Neither shows a minimum in the N region around P = 18°. In contrast, the OL24 parameterization does exhibit a minimum and lowers the N region compared to the previous *ff*s, with the energy now about 0.8 kcal/mol above the S minimum for dA and dG and about 1.2 kcal/mol for dC and dT. Note that the bsc1 *ff* provides an even less stable N region than OL15.^10^ As shown below, these changes result in a more stable A-DNA conformation with OL24. The barrier height separating the N and S regions is slightly little more than 2 kcal/mol, which is easily overcome on our simulations time scales, resulting in frequent interchanges between S and N regions (see below).

### The Percentage of N puckering in the DDD Duplex Agrees with NMR Experiments

Since DDD is one of the most extensively studied DNA duplexes and the fraction of N puckering derived from NMR measurements represents the most relevant experimental data (obtained in solution), we focus on DDD first. The percentage of N puckering for DDD, measured by 3J^17^ and RDC^18^ NMR experiments, offers a relatively consistent picture of sequence-dependent propensity for A-form in DDD. In Figure 4, these experimental data are compared with results from the OL21 and OL24 *ff*s. The OL24 sugar pucker parameterization clearly increases the population of the N region and is consistent with the NMR experiments, both in the overall percentage of N puckering and in sequence dependence. This represents a notable improvement compared to OL21, as well as the older OL15 and bsc1 *ff*s (see Ref.^10^). The development of P over time is shown in Supporting Information, Figure S1, which illustrates that interchanges between S and N puckerings occur relatively frequently, on the order of tens of ns, and thus the equilibrium distribution is reached relatively quickly in our 5 μs simulations.

**Figure 4.**
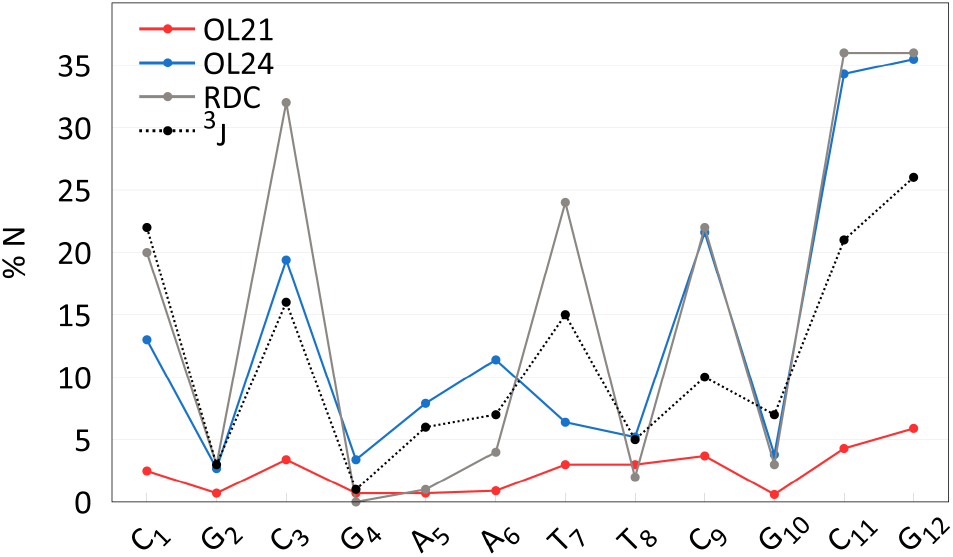
Percentage of N puckering in DDD-r simulation, averaged over 5 μs of OL24 and 2 μs of OL21 MD simulations, compared with NMR experiment.

### B-DNA Geometry Remains Virtually Unchanged by the OL24 P/χ Modification

The effect of the OL24 modification on the DDD helical parameters, groove widths, RMSD, and backbone dihedral angles is detailed in Table 3 for the DDD-r simulations (with restrained H-bonding of terminal base pairs). Overall, the geometry of DDD is only marginally altered and remains very close to that obtained with the OL21 *ff* and to experimental data. The average RMSD from the PDB structure 1BNA is 1.25 Å for OL21 and 1.30 Å for OL24 (excluding the two terminal base pairs from each end), which represents a very small increase given that we are comparing a two-state (S/N pucker) system with a static reference geometry. The RMSD remains stable throughout the simulation (Supporting Information, Figure S2). The level of fraying observed in the unrestrained DDD simulation (Supporting Information, Figure S3) is relatively low and comparable to that seen with OL15 and OL21.

**Table 3.** Helical parameters and dihedral angles characterizing DDD in MD simulations. Averages from a 5 μs OL24 DDD-r simulation are compared with those from a 2 μs OL21 simulation^13^ and averages over experimental data taken from Ref. ^12^. The averages are calculated only over the canonical (BI) regions of the dihedral potentials. Two base pairs at each end were excluded.

|  | X-RAY | OL21 | OL24 |
| --- | --- | --- | --- |
| $\alpha$/deg | 298.7 | 294.9 | 293.3 |
| $\beta$/deg | 176.6 | 180.7 | 169.7 |
| $\gamma$/deg | 52.6 | 50.2 | 51.6 |
| $\delta$/deg | 122.4 | 133.5 | 142.5 |
| $\epsilon$/deg | 180.1 | 181.3 | 189.6 |
| $\zeta$/deg | 268.6 | 266.7 | 268.5 |
| $\chi$/deg | 247.6 | 253.3 | 246.4 |
| P/deg | 130.1 | 151.7 | 150.6 |
| mg width/Å | 10.6 | 11.1 | 11.3 |
| Mg width/Å | 17.9 | 17.9 | 18.2 |
| shift/Å | 0.0 | 0.0 | 0.0 |
| slide/Å | 0.0 | 0.0 | -0.2 |
| rise/Å | 3.3 | 3.3 | 3.3 |
| tilt/deg | -0.2 | 0.1 | 0.0 |
| roll/deg | 2.0 | 1.9 | 1.4 |
| twist/deg | 33.6 | 35.2 | 35.0 |
| shear/Å | 0.0 | 0.0 | 0.0 |
| buckle/deg | 0.9 | 0.1 | 0.0 |
| stretch/Å | -0.1 | 0.0 | 0.0 |
| propeller/deg | -11.8 | -11.7 | -10.9 |
| stagger/Å | 0.1 | 0.0 | 0.1 |
| opening/deg | 2.1 | -0.2 | -0.6 |
| x-displ. /Å | -0.5 | -0.4 | -0.7 |
| y-displ. /Å | -0.1 | 0.0 | 0.0 |
| hrise/deg | 3.3 | 3.3 | 3.2 |
| incl/deg | 4.3 | 3.5 | 3.0 |
| tip/deg | 0.5 | -0.1 | 0.0 |
| htwist/deg | 34.0 | 36.1 | 36.0 |

The B-DNA backbone is characterized by a dynamic equilibrium of multiple conformations, which, in addition to A/B geometry substates, includes the BI/BII equilibrium; the BI substate (ε/ζ = t/g-) is dominant and the BII (ε/ζ = g-/t) is less populated in most base pair steps. The propensity for BII conformation is strongly sequence dependent. Figure 5A compares available experimental (NMR) data with results from OL21 and OL24 simulations. The sequence dependence of BII population is similar for both *ff*s. The total percentage of BII conformations is slightly higher with OL24 (28.2%) compared to OL21 (24.7%), with one terminal base pair excluded.

**Figure 5.**
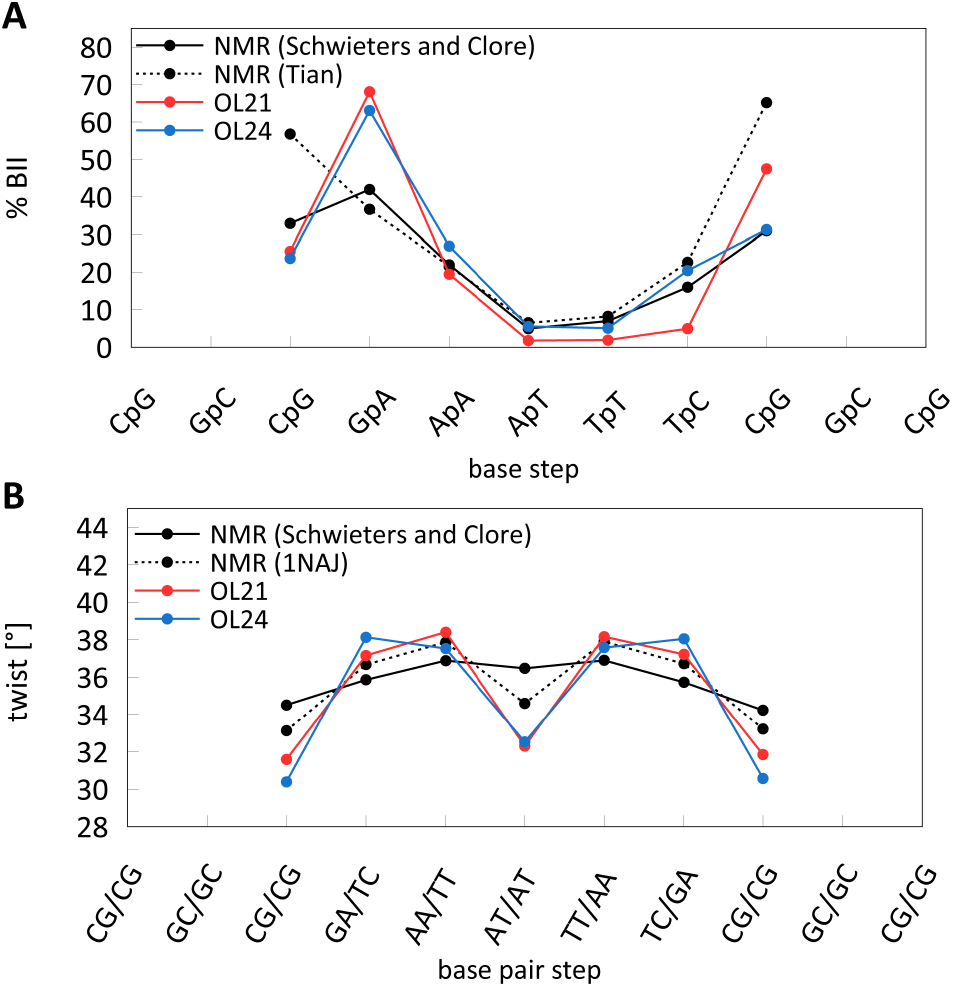
The sequence dependence of BII populations and helical twist for DDD from 5 μs OL24 DDD-r simulation compared with 2 μs OL21 simulation^13^ and NMR experiments.^66-68^

There are other minor backbone substates present in the simulations with OL24, OL21 and older *ff*s. Notably, the flips of the α/γ torsion angles, especially the α/γ = g+/t flip (compare to the canonical conformation of α/γ = g-/g+), were severely overpopulated in ff99,^55^ which necessitated the bsc0 correction.^14^ In OL21, the percentage of non-canonical α/γ substates was very low, around 1.5%. In OL24, this population increases to about 4.5%, which is still relatively low. However, unlike the excessive and spurious α/γ = g+/t conformers observed in ff99 - which were not significantly populated in X-ray structures - the conformers observed in OL24 are different. They are often associated with N puckering or changes in other backbone angles. According to the NtC annotation by černý et al.,^65^ the four main conformers found in OL24 simulations are BB05 (1.5%), BA13 (1.3%), BA16 (0.6%), and BA10 (0.3%). These conformers, described in the database study of černý et al., are part of the natural repertoire of minor B-DNA backbone conformations and may contribute to a more accurate description of interaction motifs in complexes formed by DNA.

The sequence dependence of helical parameters in OL24 is very similar to that of the previous *ff* variants (Supporting Information, Figure S4), including the helical twist, which is compared with NMR and X-ray data in Figure 5B. The distributions of backbone dihedral angles for OL24 are only slightly changed compared to OL21, except for the δ backbone dihedral angle, for which we observe a desirable increase in the N region and a small shift in the S region (Supporting Information, Figure S5).

### Other DNA Duplexes

Three DNA duplexes, for which percentages of N puckering are available from ^3^J NMR measurements,^47-49^ are compared with MD simulation results in Figure 6. For a 10-mer with mixed AT/CG content, the agreement of %N with experimental data is similarly good as observed for DDD, with average N population of 14.4% for OL24 and 10.9% from experiment.^49^ The sequence-dependence is also very similar. This represents a significant improvement over OL21, which showed only 3.2 % of N puckering, clearly too low. For both *ff*s, the helical parameters are typical of B-DNA: x-displacement is -0.7 Å for OL21 and -0.9 Å for OL24, and inclination is 3.0° for OL21 and 2.5° for OL24. The RMSD with respect to the initial NMR structure is 1.6 Å for OL21 and 1.9 Å a for OL24.

**Figure 6.**
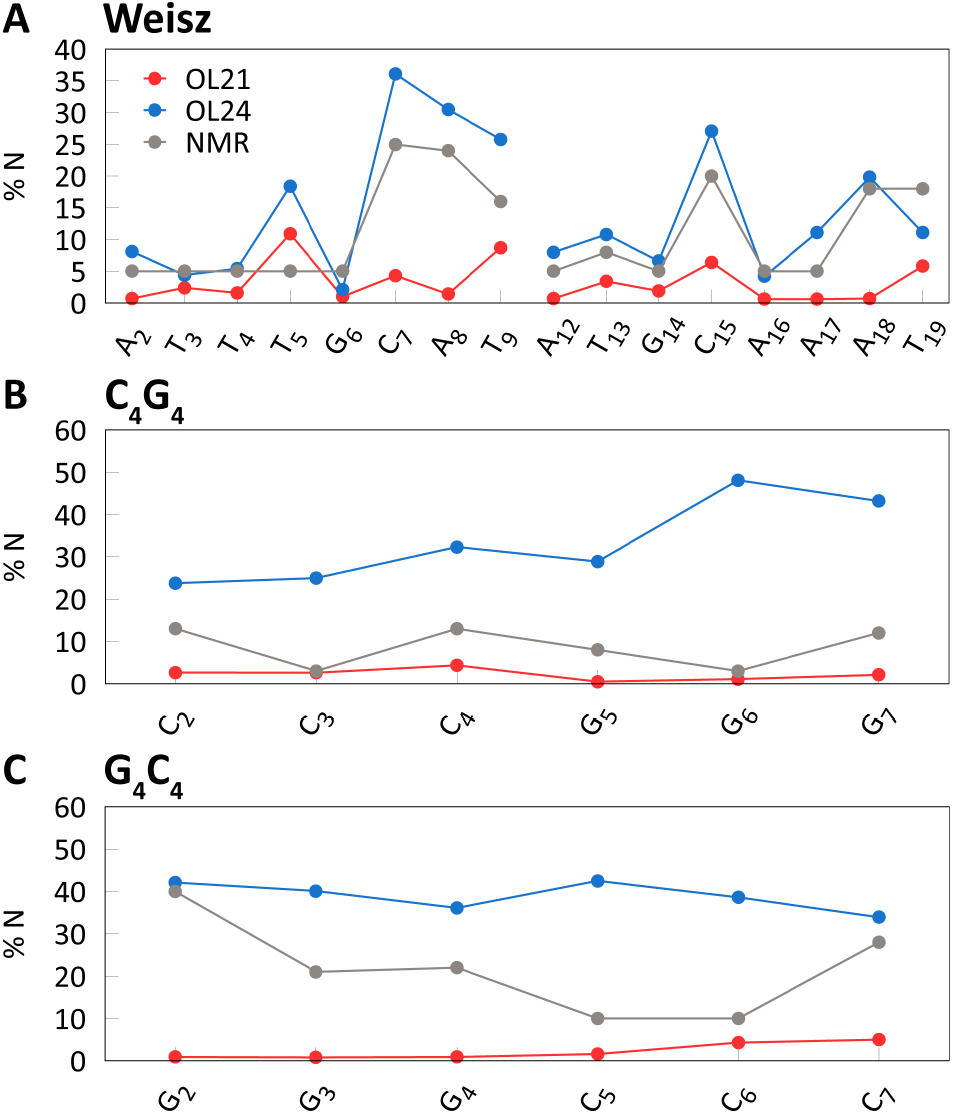
Percentages of N puckering in three DNA duplexes in aqueous solution. Averages from 2 μs simulations with one terminal base pair excluded.

In case of the pure CG sequences C_4_G_4_ and G_4_C_4_, the population of the N puckering is again underestimated by OL21 but appears to be overestimated by OL24 (Figure 6). Both C_4_G_4_ and G_4_C_4_ are unusual sequences that exhibit partly A-type base stacking in aqueous solution, as suggested by CD spectroscopy, but with predominantly B-type sugar puckering according to ^3^J NMR experiments.^47, 48^ Interestingly, a similar A-type base stacking, transitioning between A- and B-type, was observed in the crystal structure of d(CATGGGCCCATG) duplex with a G_3_C_3_ core,^69^ which features all nucleotides in the N form. The percentages of N puckering and helical parameters relevant to A/B geometries for MD simulations and experiments are compared in Table 4. Thus, it seems that this near-A-type base stacking can be consistent with sugar puckering that is both purely N or predominantly S.

**Table 4.** RMSD, inclination, x-displacement, and percentage of N puckering (%N) for three DNA duplexes in aqueous solution. Averages from 2 μs simulations, terminal base pairs were excluded.

|  |  | inclination [°] | x-displ. [Å] | % N |
| --- | --- | --- | --- | --- |
| B-DNA |  | 1.5 | 0 |  |
| A-DNA |  | 20.7 | -5.3 |  |
| C<sub>4</sub>G<sub>4</sub> | OL21 | 6.4 | -0.5 | 2.2 |
|  | OL24 | 8.2 | -2.1 | 33.5 |
|  | exp<sup>47</sup> |  |  | 8.7 |
| G<sub>4</sub>C<sub>4</sub> | OL21 | 5.3 | -0.5 | 2.3 |
|  | OL24 | 8.4 | -2.6 | 38.9 |
|  | exp<sup>48</sup> |  |  | 21.8 |
| G<sub>3</sub>C<sub>3</sub> | exp<sup>69</sup> | 5.5 | -2.9 | 100.0 |

Although %N appears to be overestimated with OL24 for the C_4_G_4_ and G_4_C_4_ duplexes, the helical parameters are much closer to the intermediate A/B form discussed above (Table 4). While OL21 predicts helical parameters characteristic of a purely B-type stacking, OL24 seems to describe the geometry of the C_4_G_4_ and G_4_C_4_ molecules more accurately.

### Hybrid DNA/RNA Duplexes Show Higher %N in the DNA Strand with OL24

Two DNA/RNA hybrids were investigated, one with a predominantly purine DNA strand and the other with a predominantly pyrimidine DNA strand (PDB IDs 1DRR and 1RRD, respectively;^27^ Figure 7). In a hybrid, the RNA molecule tends to push the DNA strand closer to the A-form, as suggested by NMR experiment,^27^ as well as X-ray structures, where the deoxyribose puckering is frequently N.^70, 71^ The overall helical geometry of the hybrids in solution is intermediate between A-to B-forms (Figure 8), as also observed in MD simulations.^19, 72, 73^

**Figure 7.**
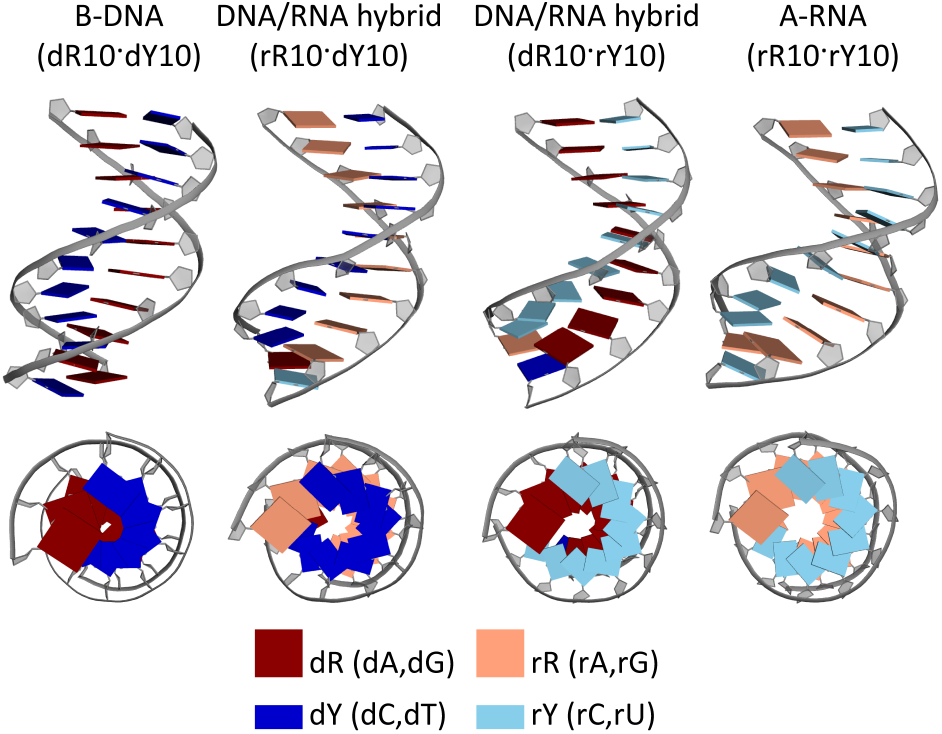
Comparison of NMR structures of DNA/RNA hybrid geometries with ideal B-DNA and A-RNA structures, taken from Ref^27^ (PDB IDs 1AXP, 1RRD, 1DRR and 1RRR, from left to right). Note the ‘hole’ in the center of the duplexes and compare with x-displacement values from simulations for these duplexes, as shown in Table 4.

**Figure 8.**
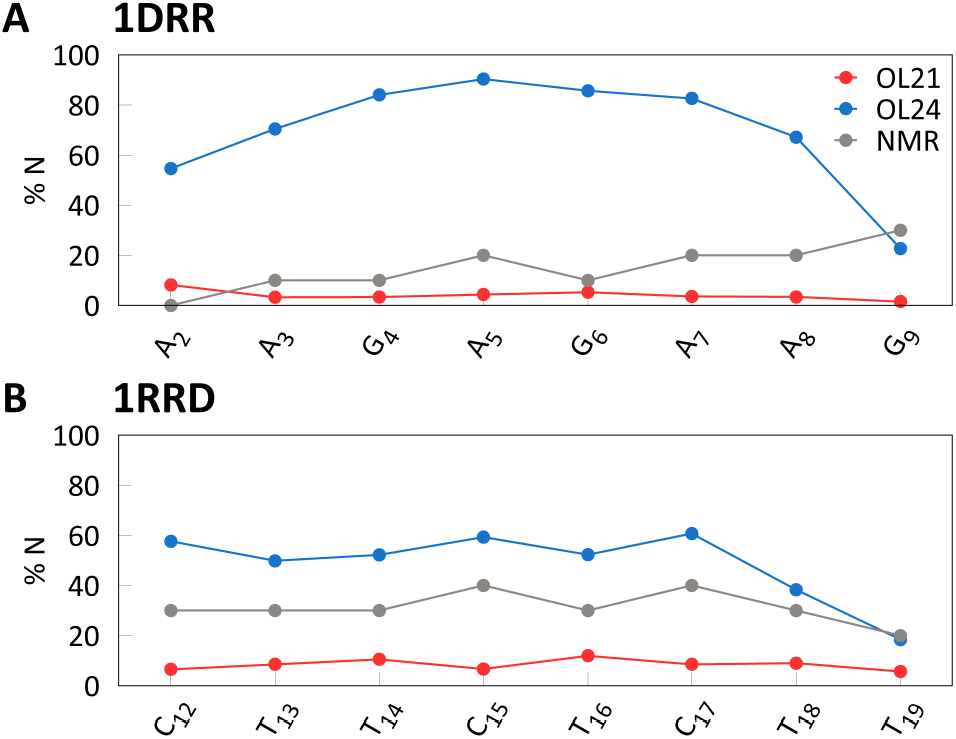
Sequence dependence of the percentage of N pucker (%N) in the DNA strand in simulations of DNA/RNA hybrids compared with experiment.^27^

While it was suggested that this intermediate form features O4’-endo sugar puckering,^74^ the prevailing opinion is that it features a mixture of N and S sugar puckering,^27, 72, 75^ with the population of the N puckering in the DNA strand increasing when compared to a B-DNA double helix. In X-ray structures, and sometimes also in NMR structures, all DNA nucleotides may be found in the N conformation.^70, 76^

Figure 8 compares the populations of the N state in the DNA strand of hybrids with NMR experimental data. While the OL21 *ff* predicts populations of N puckering that are too low, OL24 seems to overestimate %N in both sequences. However, the %N values from NMR data were obtained as averages over an ensemble of 10 NMR structures,^27^ which may introduce inaccuracies. It is also worth noting that much higher %N content has been reported for other hybrids in both X-ray^70^ and NMR structures.^76^ Therefore, we suggest that OL24 provides a better description of N pucker population in DNA/RNA hybrid duplexes than the older AMBER *ff*s.

Helical parameters discriminating between A- and B-form are shown in Table 5, along with RMSD values relative to the first structure in the NMR ensemble and average %N. In terms of helical parameters, both structures simulated with OL24 fall between the A- and B-forms and are shifted closer to the A-form compared to OL21. This is desirable, as current AMBER *ff*s have been shown to predict helical parameters of hybrids that are somewhat biased towards the B-form.^19^ Therefore, OL24 appears to provide an improvement.

**Table 5.** DNA/RNA hybrids with predominantly purine (1DDR) and pyrimidine (1RRD) DNA strands. Average over 2 μs simulations, DNA strand only; terminal base pairs were excluded.

|  |  | incl [°] | x-disp [Å] | % N | RMSD [Å] |
| --- | --- | --- | --- | --- | --- |
| 1DDR | OL21 | 15.7 | -2.9 | 4.1 | 2.1 |
|  | OL24 | 16.3 | -4.2 | 69.7 | 1.7 |
|  | expt.<sup>27</sup> | 8.0 | -4.65 | 15 |  |
| 1RRD | OL21 | 7.3 | -3.1 | 8.4 | 1.2 |
|  | OL24 | 8.8 | -3.6 | 48.6 | 1.3 |
|  | expt.<sup>27</sup> | 8.0 | -4.2 | 31 |  |
| B-DNA<sup>77, 78</sup> |  | 1.5 | 0.0 |  |  |
| A-DNA<sup>77, 78</sup> |  | 20.7 | -5.3 |  |  |

### B-DNA Transitions to A-DNA in Ethanol Solutions with OL24

In concentrated ethanolic solutions, the DNA duplex is known to undergo a transition from B-to A-DNA form.^16^ Since the ethanol concentration required for this transition varies depending on the sequence - with GC-rich DNAs requiring lower concentrations - we chose a relatively high (85%) ethanol concentration to ensure that the DDD sequence can adopt the A-form. In our previous works, none of the current AMBER *ff*s (OL15, bsc0 and bsc1) predicted stable A-DNA - when starting from the B-form, the duplex largely remained in the B-form, and when starting from the A-form, the duplex quickly transitioned to B-DNA.

Here, we perform these simulations with OL21 and OL24 *ff*s. The progress of the transition, shown in Figure 9, is monitored by inclination and x-displacement, which we suggested as better indicators of the A/B form than RMSD.^10^ Additionally, we also present RMSD relative to the idealized A-DNA from the NAB.^45, 46^

**Figure 9.**
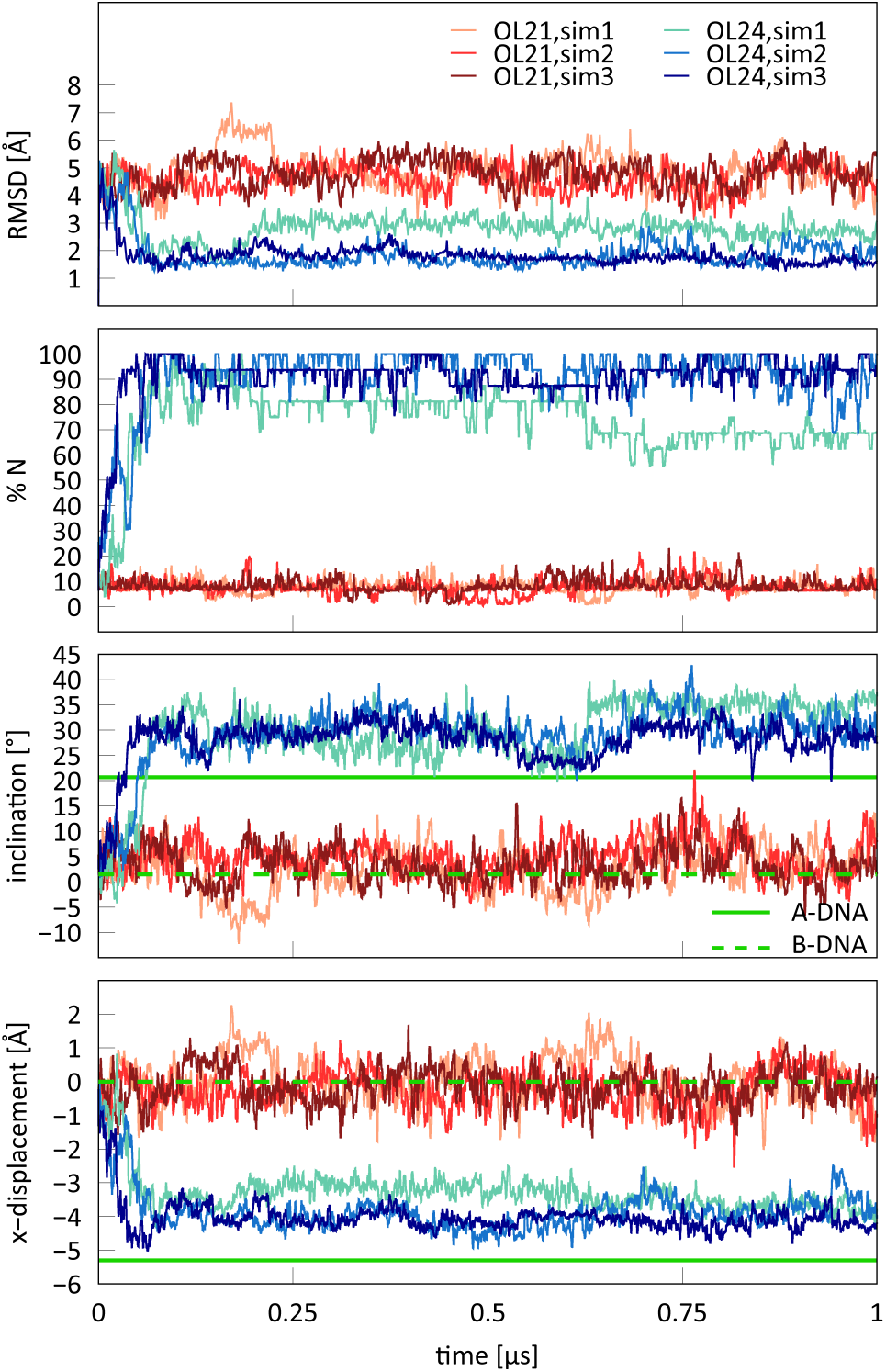
Three independent OL21 and OL24 simulations of DDD starting from B-DNA conformation in 85% ethanol solution. Only the OL24 *ff* models the B-to-A transition. Reference values for inclination and x-displacement of A- and B-DNA forms are the same as in Table 4. RMSD is relative to the idealized A-form.

**Figure 10.**
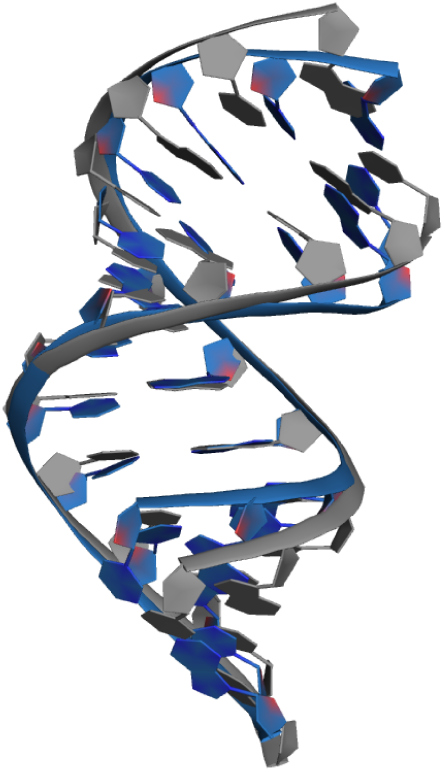
Overlay of the ideal A-DNA structure (gray, obtained from NAB) and the average simulated structure (OL24_sim2, second half of the simulation) in ethanol/water solution (blue), illustrating the narrowing of the major groove in the MD simulation.

When starting from the B-DNA form, all three independent OL24 simulations transitioned from B-to A-DNA form relatively quickly, within about 100 ns (Figure 9). The OL21 simulations remained in the B-form. This indicates that the A/B equilibrium in concentrated ethanol solutions is much better described by OL24 than by the OL21 and OL15 *ff*s, where no B-to A-transition was observed. Note that the bsc1 *ff* was also unable to model a stable A-DNA form.^10^

We should note that in one of the three simulations (OL24_sim1), we observed an increased RMSD and a broken base pair near the middle of the structure. However, the remaining bases retained the A-DNA form. Although such an isolated break in the A-DNA helical structure may not be detected by CD spectroscopy, and the simulated structure could still be in agreement with the experiment, we suggest that this anomaly is likely an artefact of the simulation rather than a genuine behavior of A-DNA structure in ethanol solution. We noticed a narrow major groove () with strongly bound cations inside, which we suspect is a consequence of imbalances in the description of the cation/DNA/water+ethanol equilibrium. It is important to note that the parameters for the ethanol are relatively old and were not derived with consideration for the solvation energies of monovalent cations in water mixtures. Nevertheless, despite the somewhat deformed A-DNA structure in one of the simulations, the average helical parameters and fraction of N puckering (accompanied with χ = *anti*) clearly indicate a transition to the A-form.

### OL24 Stabilizes the A-form in Protein-DNA Complexes

In our previous work, we have shown that A-form of DNA is generally not stable enough in simulations of eight protein-DNA complexes using the current AMBER *ff*s OL15 and bsc1.^11^ Here, we focus on two of those complexes that were characterized by a high A-DNA content: TnpA transposase (PDB ID 2VJV)^43^ and PvuII endonuclease (PDB ID 3PVI),^44^ shown in Figure 11. Figures 12 and 13 show the average percentage of N sugar puckering during the MD simulations, but only for residues that were in the A-form in the crystal structure. As discussed previously, the fraction of A form quickly decreases in simulations with OL15 and bsc1 *ff*s,^11^ and simulations with the OL21 *ff* show the same trend. In the case of the PvuII endonuclease (3PVI), disappearance of the A-DNA conformation is accompanied by a significant increase of RMSD of the DNA helix and the whole complex (Figure 12). In case of TnpA transposase, the increase in RMSD is not noticeable, possibly because there is a smaller fraction of A-like nucleotides in the initial X-ray structure.

**Figure 11.**
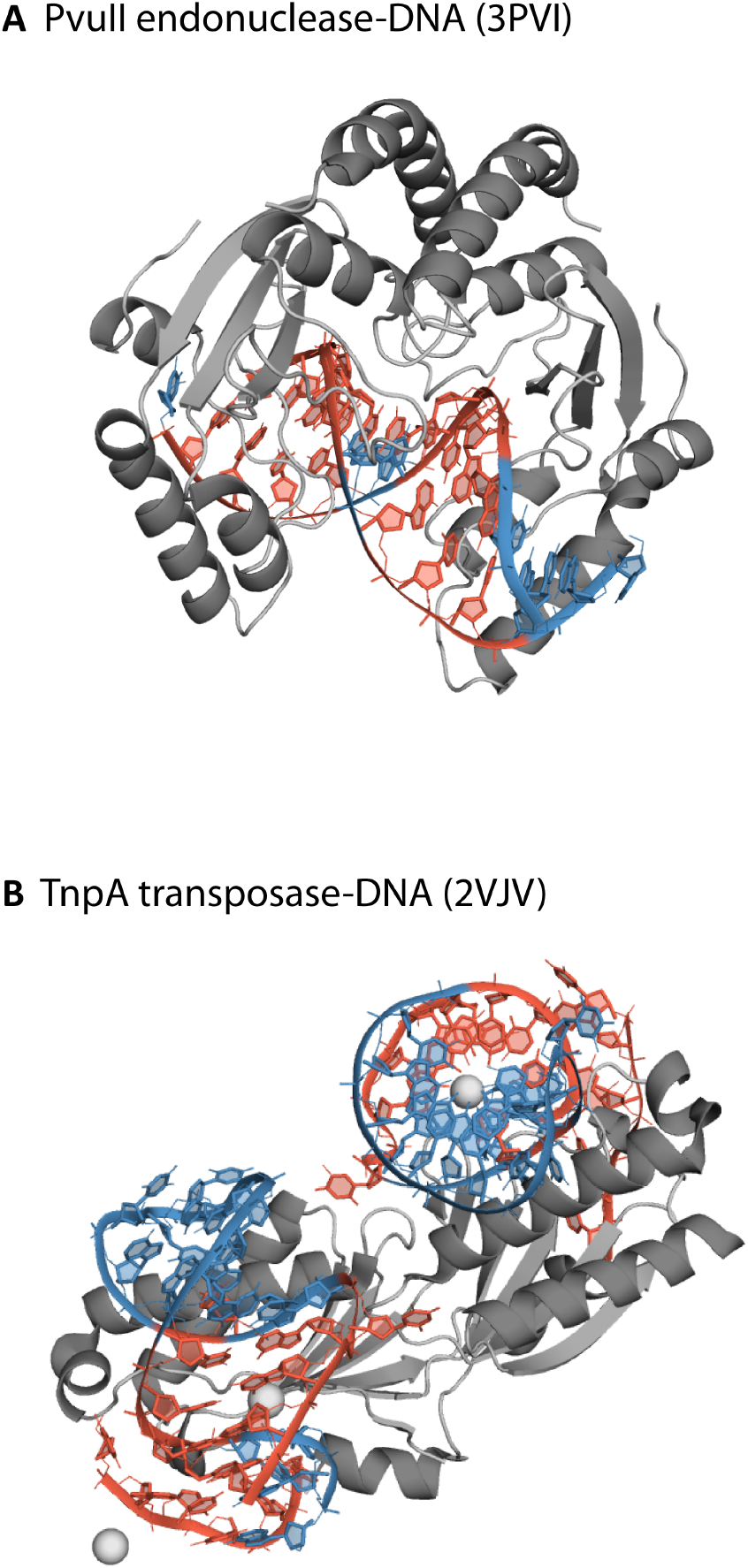
Simulated protein-DNA complexes. Residues shown in red are in the A-form in the X-ray structure.

**Figure 12.**
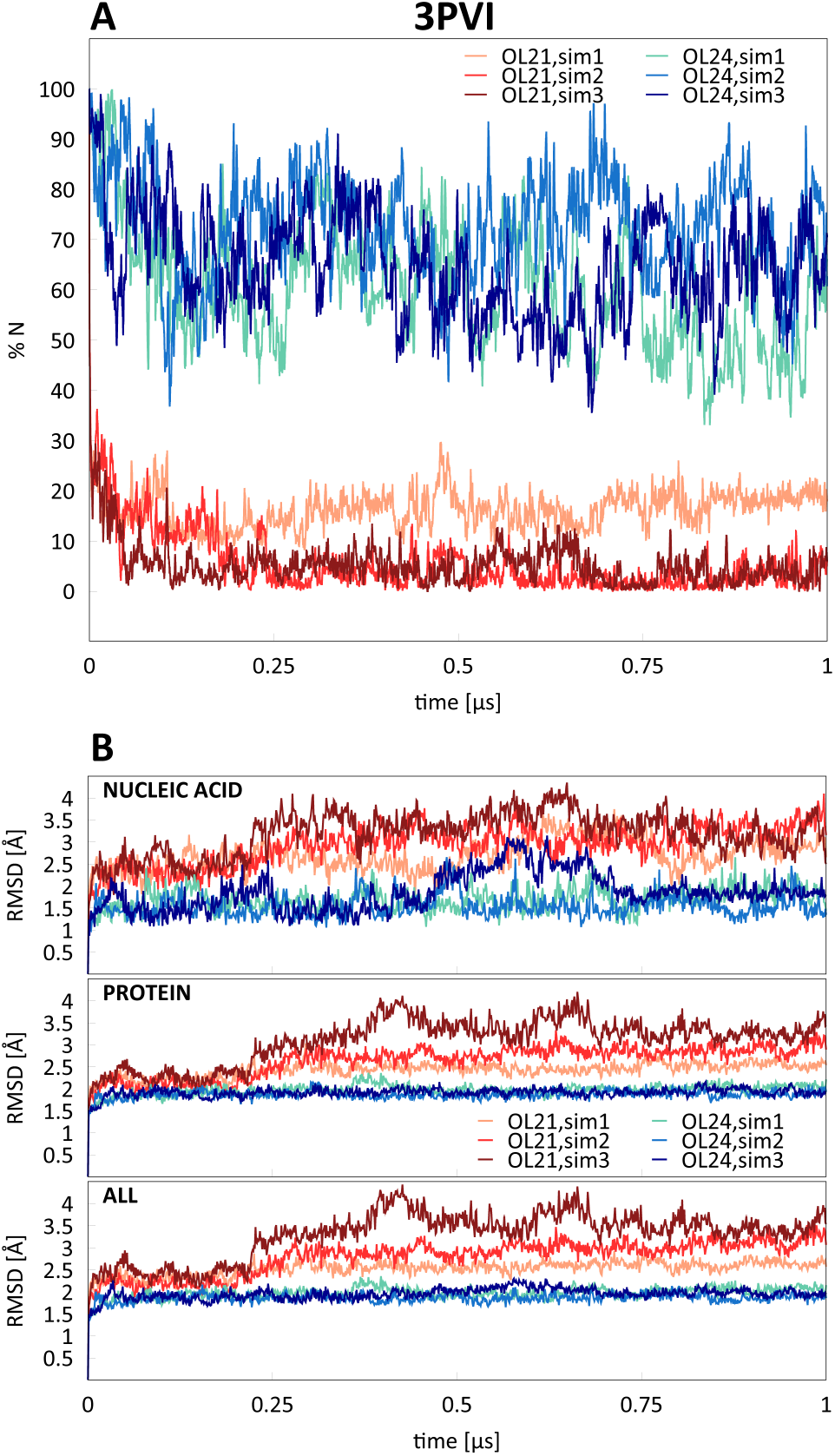
Average percentages of N sugar pucker (%N) in residues that adopt the A-form in the starting X-ray structure and RMSD for three independent simulations of PvuII endonuclease (3PVI) with OL21 and OL24 *ff*s.

**Figure 13.**
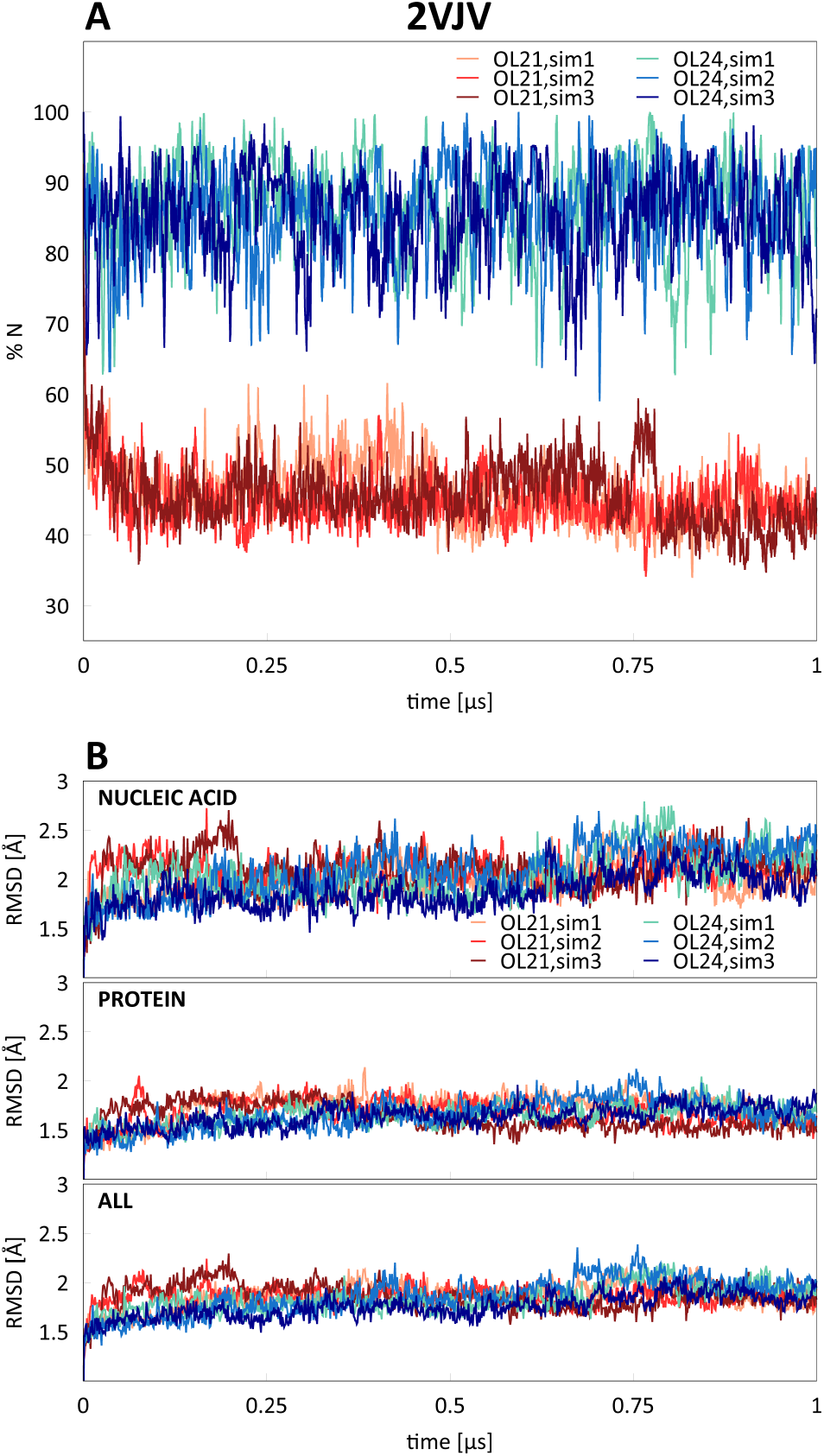
A) Average percentages of N sugar pucker (%N) in residues that adopt the A-DNA form in the starting X-ray structure and B) RMSD for three independent simulations of TnpA transposase (2VJV) with OL21 and OL24 *ff*s.

In contrast, OL24 simulations show a much higher preservation of %N for both simulated complexes. Notably, in case of the PvuII endonuclease (3PVI), this is accompanied by a significantly smaller RMSD relative to the initial X-ray structure. However, it is important to note that comparing MD simulations with X-ray structures is not straightforward. Specifically, the assumption that an A-state observed in an X-ray structure will be preserved throughout an MD simulation may be an oversimplification. X-ray structures are measured at low temperatures, where individual nucleotides are pushed to their enthalpic minima. In solution, however, these same nucleotides could exhibit equilibrium between A and B forms. Consistent with this, MD simulations often show A/B equilibria, and we can only expect that residues existing in the A conformation in the X-ray structure will show higher A-DNA content in simulations. Therefore, the relatively high %N content in the selected residues suggests that our OL24 simulations are in better agreement with X-ray experiments than the older AMBER *ff* variants.

Simulations with older *ff*s, where RMSD notably increases, are likely of only limited use and cannot be fully trusted. However, it should be noted that even when RMSD is not excessively large, the older *ff*s still predict too low stability for the A-form in protein-DNA complexes. Thus, even if geometry (judged by RMSD) of a complex appears acceptable, the thermodynamic predictions and structural details are likely to be biased when using current AMBER *ff*s. Therefore, applying the refined OL24 *ff* is advisable for simulations of protein-DNA complexes.

## Conclusions

We present a reparameterization of the deoxyribose dihedral parameters, OL24, that improves the modeling of the A/B-DNA equilibrium by stabilizing the N-puckering of the deoxyribose ring. Because it is used in conjunction with the OL3 glycosidic angle parameterization, both RNA and DNA Olomouc *ff*s now share the same glycosidic dihedral parameters. OL24 parameter files are available on our web site (ffol.upol.cz) and dihedral parameters are given in Supporting Information, Table S1.

OL24 provides results that are more consistent with available experimental data than the previous AMBER parameterizations, such as OL15, OL21, and bsc1. Specifically, OL24 predicts significantly higher populations of N puckering in DDD duplex in aqueous solution, in agreement with ^3^J and RDC NMR data. Similar increases are observed in other DNA duplexes. Unlike previous AMBER parameterizations, OL24 correctly predicts the transition from B-DNA to A-DNA in concentrated ethanol solutions, consistently stabilizing the A-type structure with high fraction of N puckering.

Importantly, OL24 maintains a highly accurate description of the overall B-DNA structure, including helical parameters, groove widths, RMSD, backbone dihedral angle distributions and BI/BII populations, with only a minimal admixture of non-canonical backbone substates. Therefore, OL24 is recommended for simulations of canonical B-DNA structures.

The OL24 modification improves the modeling of biologically relevant DNA complexes, such as DNA/RNA hybrid duplexes and protein-DNA complexes. Whereas previous AMBER parameterizations like OL15, OL21 and bsc1 strongly underestimated population of N sugar puckering in DNA/RNA hybrids, OL24 shows a much higher N state content, consistent with NMR and X-ray experiments.

For protein-DNA complexes, OL24 stabilizes the A-DNA conformation more effectively than older AMBER *ff* variants, providing a more realistic representation of protein-DNA interactions. This results in more stable protein-DNA complexes and can significantly reduce the RMSD of the simulated structures.

The improved modeling of the A/B equilibrium allows for more realistic modeling of the DNA double helix and its interactions with other biomolecules, including DNA/RNA hybrids and protein-DNA complexes, which are fundamental to molecular biology.

## Acknowledgements

This work was supported by grant 17-16107S (P.J., M.Z.) from the Grant Agency of the Czech Republic.

## Supporting Information

Details of DDD simulations: pucker time series, RMSD time series, fraying, sequence dependence of helical parameters, distributions of backbone dihedral angles and OL24 dihedral parameters.

