## Supporting Information for "Refinement of the Sugar Puckering Torsion Potential in the AMBER DNA Force Field"

**Figure S1**. Time series of sugar pucker in the DDD-r duplex simulation. One base pair at each end was excluded.

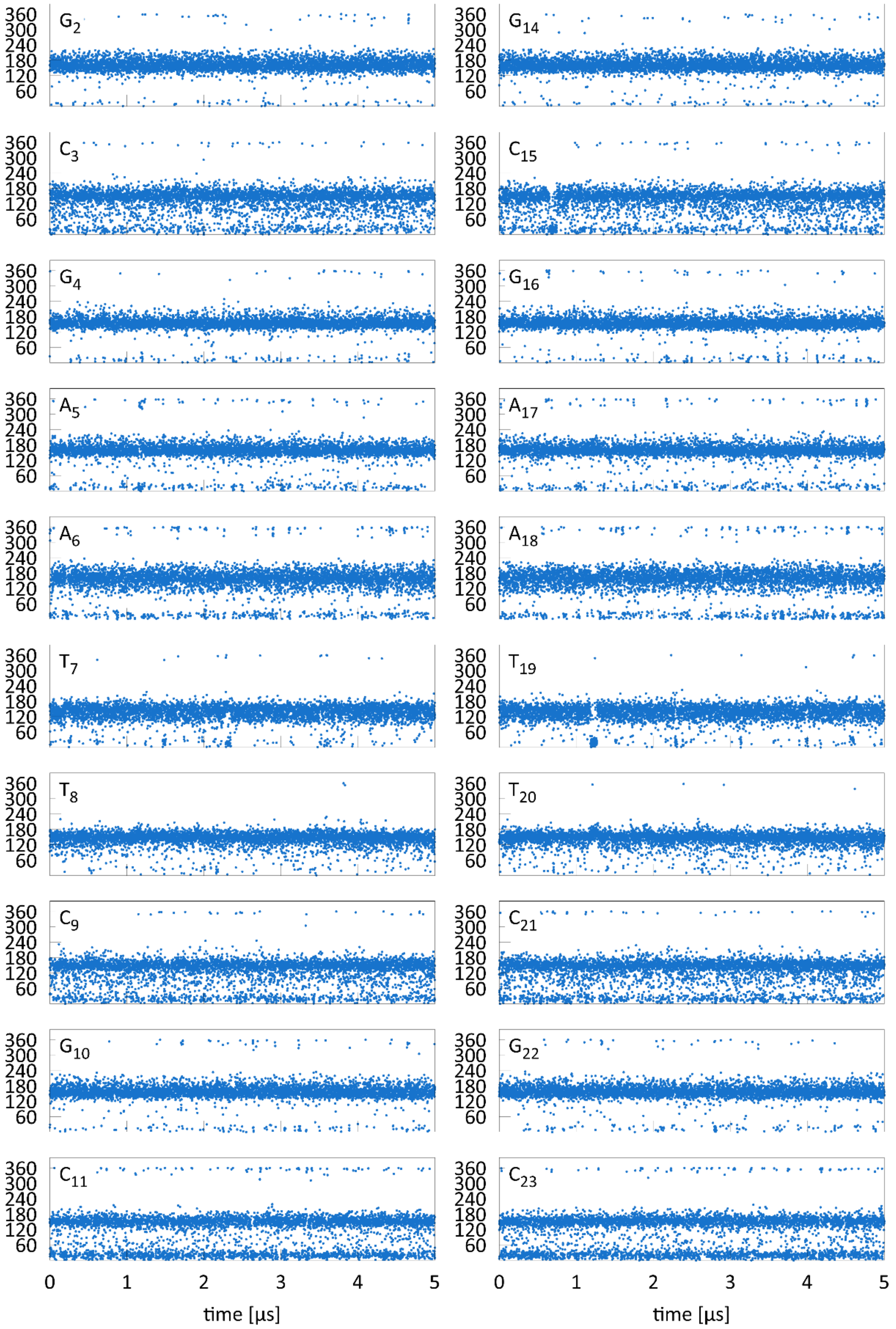

**Figure S2.** RMSD of simulated DDD relative to the 1BNA structure. Two base pairs at each end were excluded. The inset compares 5 μs OL24 simulation with 2 μs OL21 simulation.

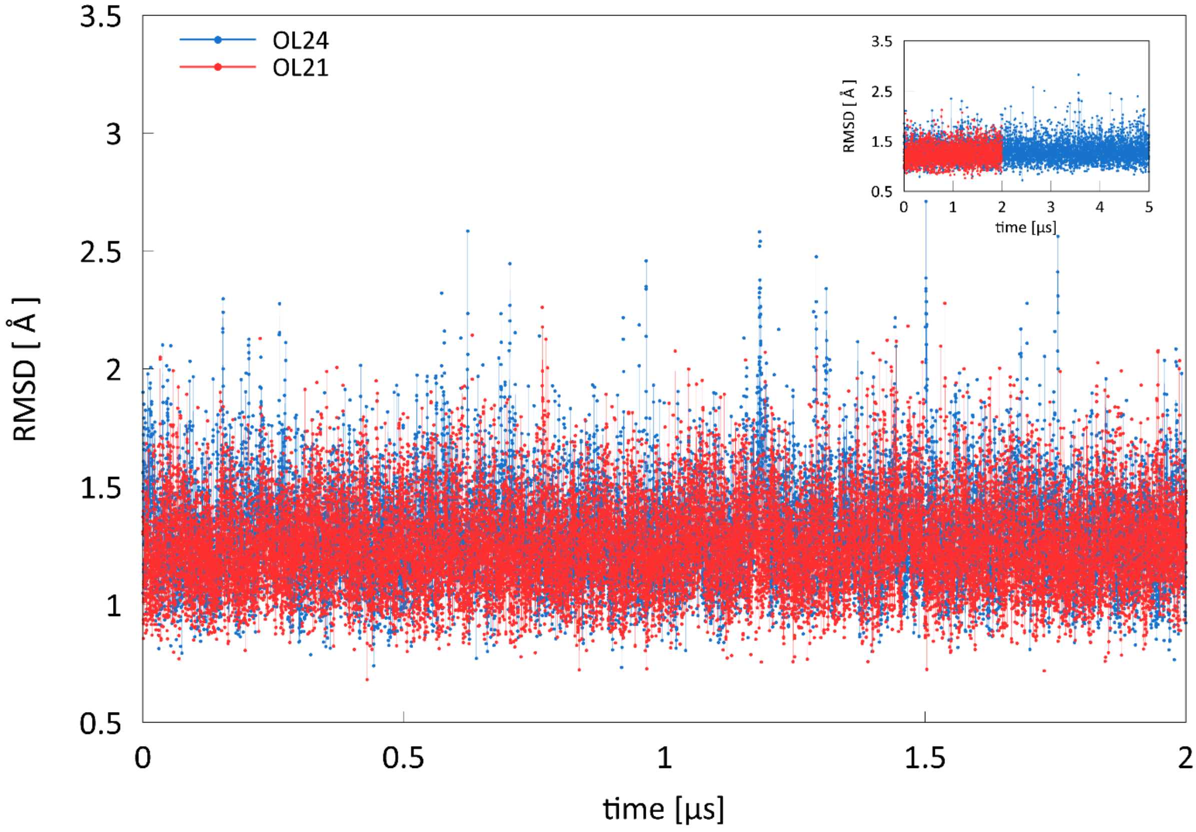

**Figure S3.** Fraying in unrestrained DDD dodecamer simulation: RMSD of the terminal bases $C_1$,$G_{24}$ and $G_{12}$,$C_{13}$.

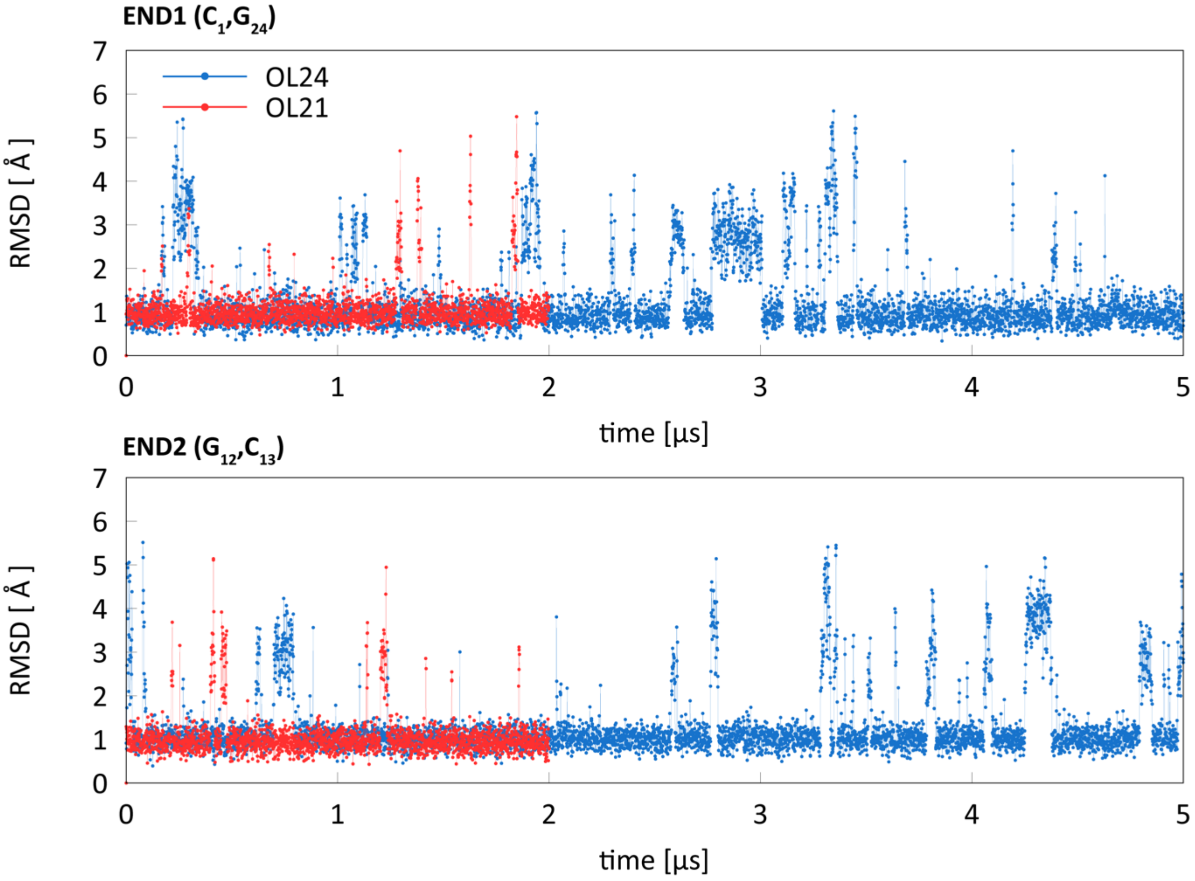

**Figure S4.** Sequence dependence of helical parameters in DDD-r simulations.

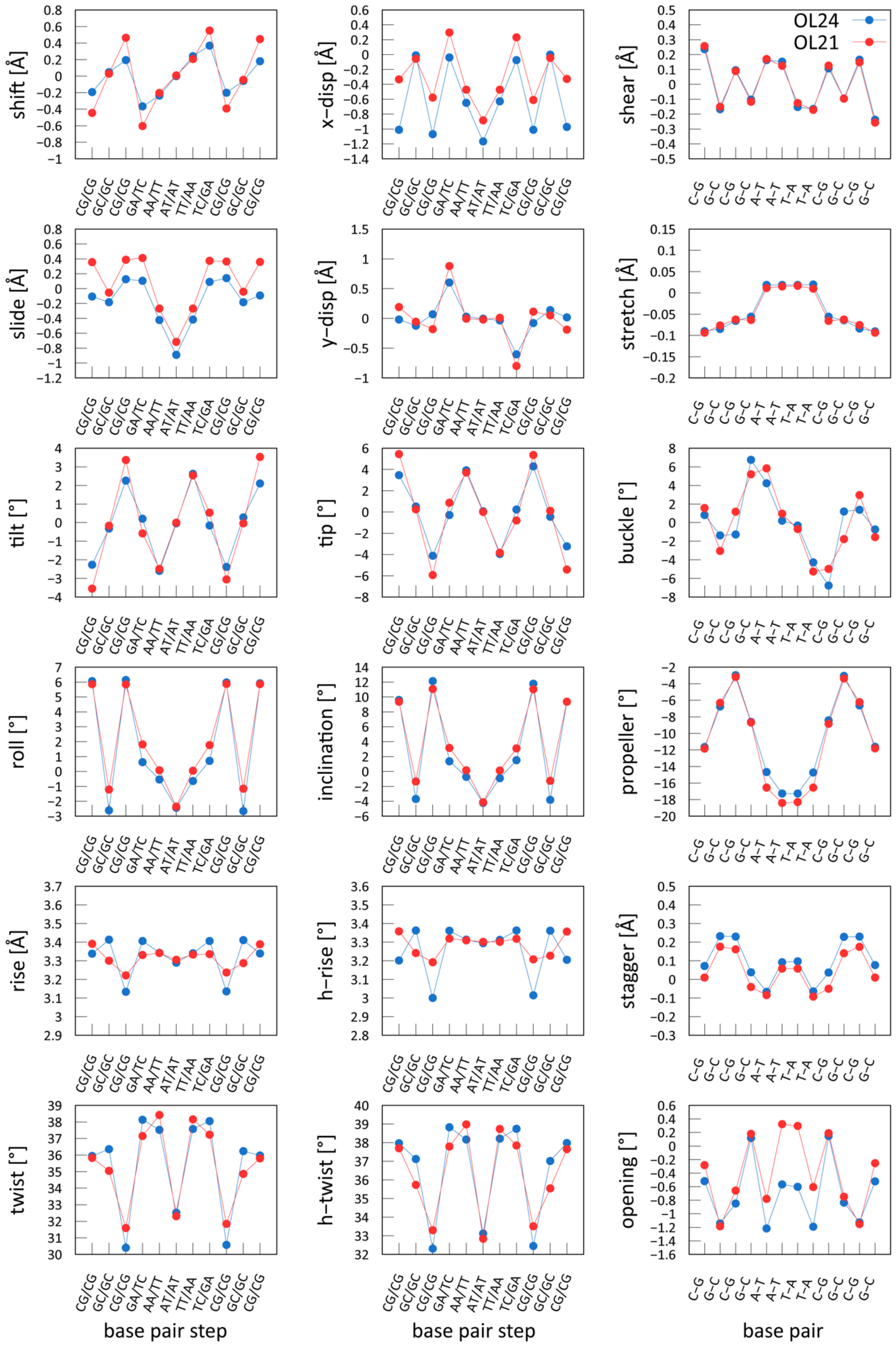

**Figure S5.** Dihedral angle distributions for DDD-r simulations. One base pair at each end was excluded.

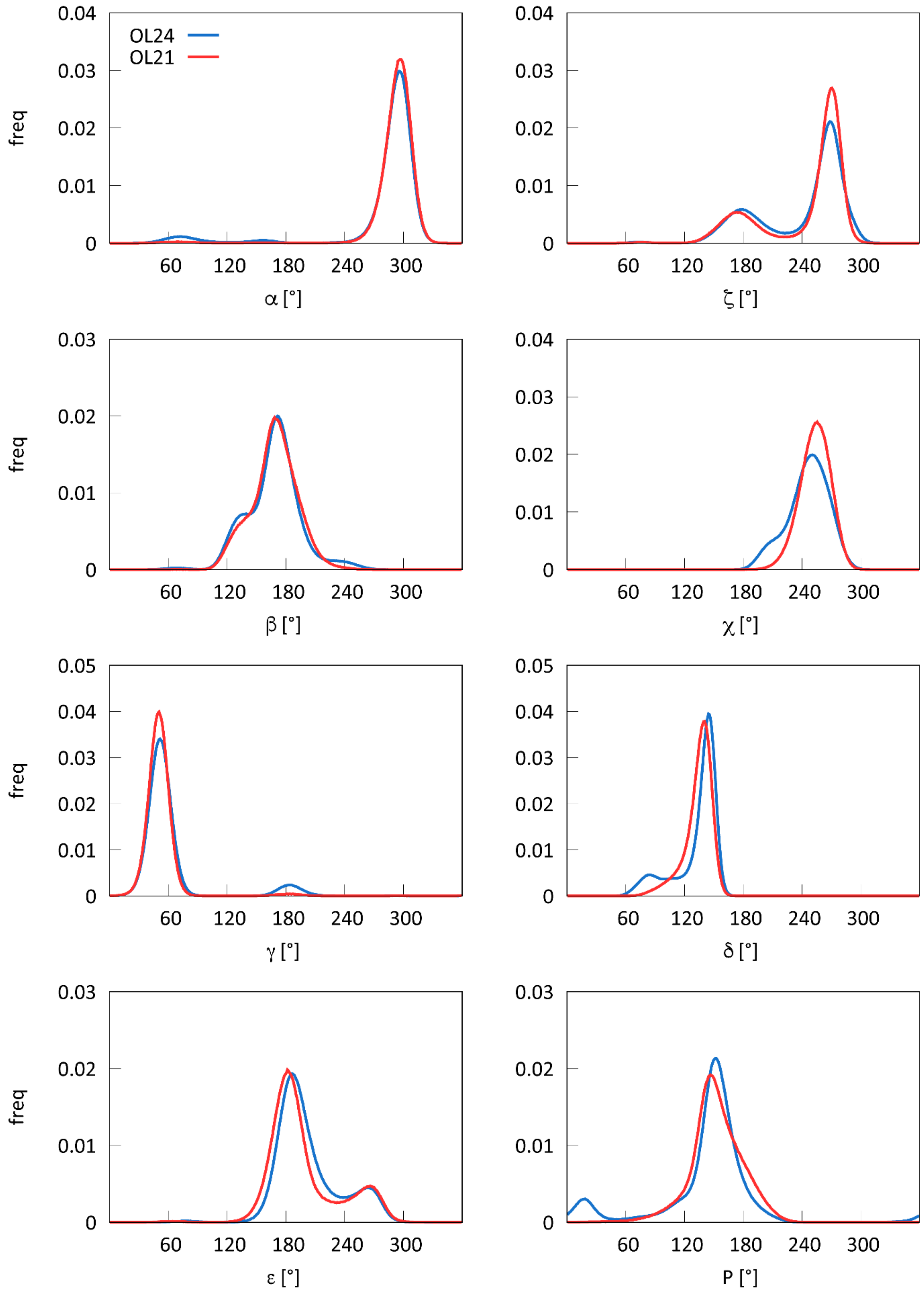

**Table S1.** OL24 deoxyribose dihedral angle parameters.

available from ffol.upol.cz
